# Dynamics of fibril collagen remodeling by tumor cells using individual cell-based mathematical modeling

**DOI:** 10.1101/2021.09.21.461296

**Authors:** Sharan Poonja, Mehdi Damaghi, Katarzyna A. Rejniak

**Affiliations:** Integrated Mathematical Oncology, H. Lee Moffitt Cancer Center & Research Institute, FL; Department of Cancer Physiology, H. Lee Moffitt Cancer Center & Research Institute, FL; Oncologic Sciences Department, Morsani School of Medicine, University of South Florida, FL

## Abstract

Many solid tumors are characterized by dense extracellular matrix (ECM) composed of various ECM fibril proteins that provide structural support and biological context for the residing cells. The growing tumor cell colonies are capable of remodeling the ECM structure in tumor immediate vicinity to form specific microenvironmental niches. The changes in fibril patterns of the collagen (one of the ECM proteins) surrounding the tumor can be visualized experimentally using both histology and fluorescent imaging. In particular, three diverse tumor associated collagen signatures (TACS) were identified and related to tumor behavior, such as benign growth or invasion. Here we will use an off-lattice hybrid agent-based model (*MultiCell-LF*) to identify the rules of cell-ECM interactions that guide the development of various patterns of alignment of the ECM fibrils.

## 1. Introduction

The *in vivo* tumor microenvironments are complex and dynamically changing. The extracellular matrix that fills the space between tumor and stromal cells, is composed of approximately 300 different proteins (1, 2). The most abundant belong to a class of fibrous proteins (such as collagens, fibronectins, elastins, or laminins) that are assembled into well-organized meshes and form structural support for the residing cells (3). The ECM proteins are locally secreted by stromal cells (such as fibroblasts) and their excessive deposition (fibrosis) is often a sign of aggressive tumors (3, 4). The ECM structure can also be remodeled by tumor cells that has been associated with tumor invasiveness, metastatic spread, and increased mortality in patients with breast cancers (5, 6). The most studied ECM alteration in the tumor tissue is collagen deposition that can be visualized using the second harmonic generation (SGH) microscopy (5, 7). Collagen density and alignment can regulate cancer cell signaling, proliferation, polarity, and migration (8, 9).

In several experimental studies of breast cancer development in mice, the specific tumor-associated collagen signatures (TACS) were observed (7, 10, 11). The TACS-1 signature was detected in the areas located farther from the growing tumor cell colony and was described as unorganized fibrils with wavy appearance (compare Figure 3 in (5), Figure 1 in (10), and Figures 4ab,5a in (11)). The TACS-2 signature was characterized by stretched collagen fibrils aligned parallel to the edge of the tumor cluster (compare Figure 3 in (5), Figure 1 in (10), and Figures 4de in (11)). The TACS-3 signature was identified as collagen fibrils oriented radially from the tumor cluster, often at the site of local invasion (compare Figure 3 in (5), Figures 1, 2b in (10), and Figures 4f, 5b in (11)).

**Figure 1.**
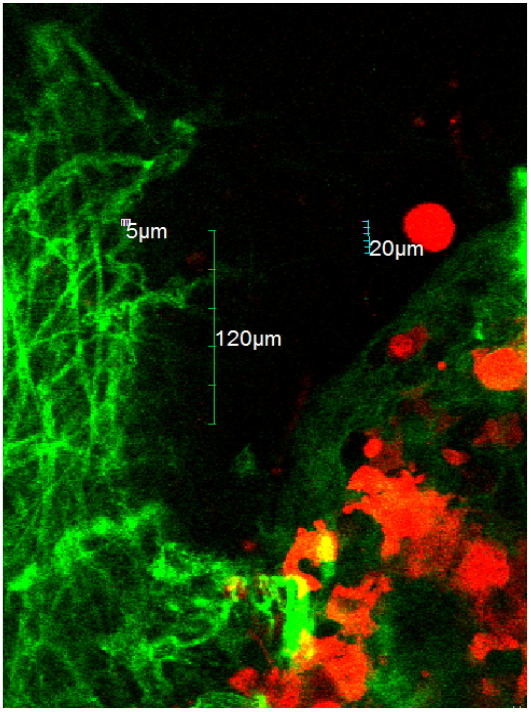
In vitro experiment showing RFP-tagged tumor cells (red) and GFP-labelled ECM collagen fibers (green).

**Figure 2.**
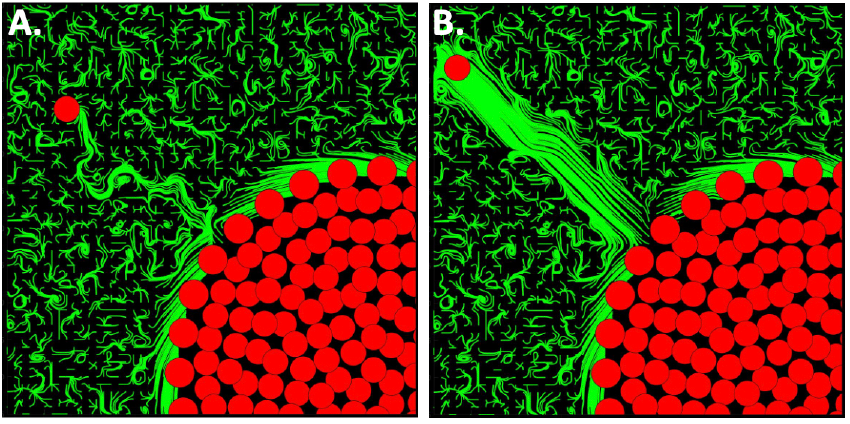
Cells migrating along the randomly aligned fibrils with ECM remodeling: **B**. a cell with low persistence migration *β* = 0.1 and ECM with low compliance *α* = 0.4. **C**. a cell with high persistence migration *β* = 0.9 and high ECM compliance *α* = 0.9.

**Figure 3.**
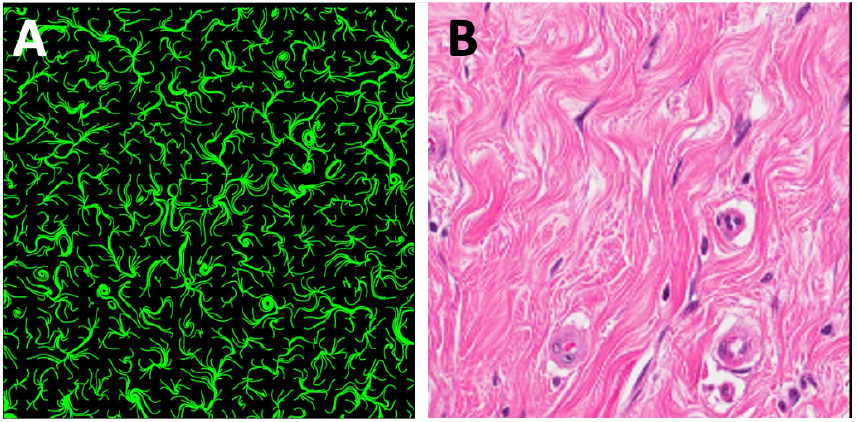
Model of TACS-1. **A**. Simulated ECM with randomly aligned fibrils and **B**. the corresponding wavy ECM pattern from histology of a normal mammary tissue.

**Figure 4.**
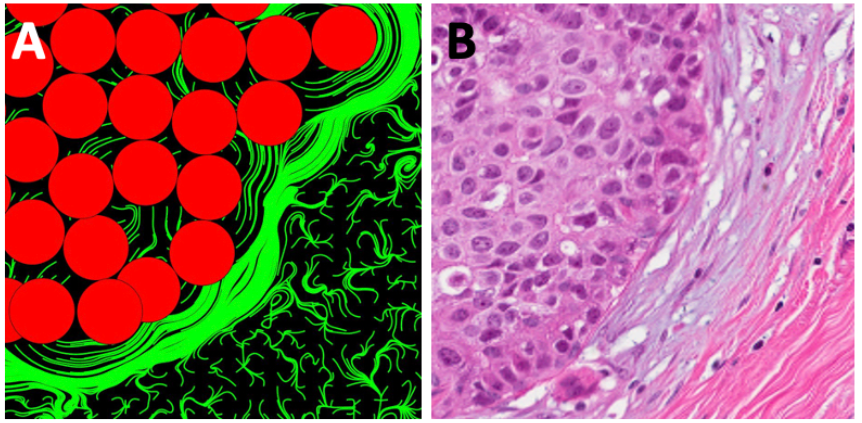
Model of TACS-2. **A**. Simulated ECM with fibrils aligned parallel to the growing cell cluster and **B**. the corresponding ECM patterns along the boundary of the DCIS

**Figure 5.**
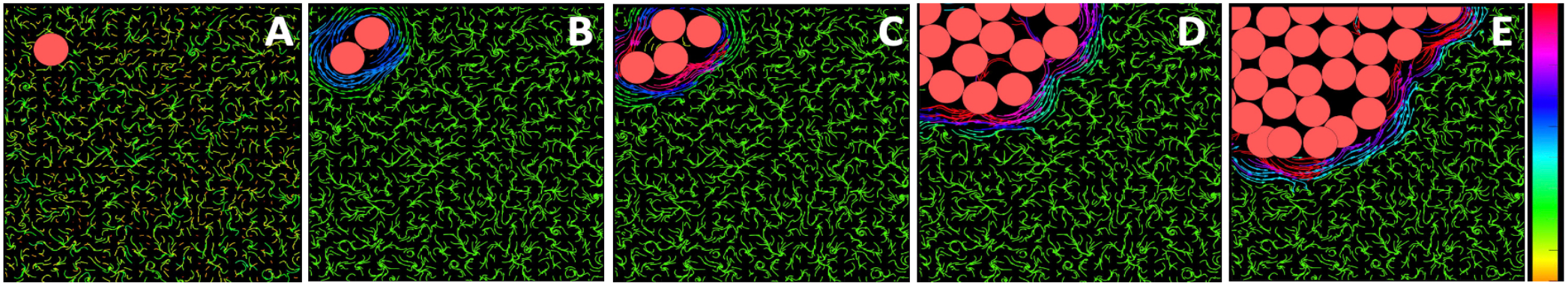
Development of TACS-2 in the mathematical model. **A-E**. Snapshots from a simulation showing the emergence of stiff ECM fibrils aligned perpendicular to a growing cell cluster. The local ECM fibril stiffness is shown in a color scale.

Such changes in ECM structure and cell-ECM interactions can only be captured by individual cell-based models (called also agent-based models, ABMs). These models are capable of reproducing various tumor morphologies, growth dynamics and tissue heterogeneities (12, 13). In particular, the off-lattice models that we and others developed (14-16) can also incorporate cell and ECM mechanics, and diverse tissue architectures, such as mammary ducts or multicellular organoids. Our *MultiCell-LF* (multi-cellular lattice-free) model (16-20) can trace physical interactions between individual cells (*i*.*e*., repulsive or adhesive forces), between cells and ECM fibrils (bundle formation or change in fibril orientation), and the kinetics of diffusible factors (*i*.*e*., oxygen or enzymes) that the cells and ECM fibrils are exposed to. Here, we will use this model for a comprehensive analysis of the emerging patterns of ECM during tumor progression.

## 2. Mathematical Model

Our *MultiCell-LF* mathematical model follows experimental data setting (Figure 1) and includes tumor cells modeled as individual agents, and ECM fibrils represented by a discrete vector field with specified fibril direction and stiffness value for each fibril. The physical interactions between both cells and fibrils are modeled using spring forces.

Each tumor cell is represented by its nucleus ***X***_*i*_ and radius *R*. To ensure that the cells did not overlap with one another, repulsive forces were applied to all cells. Let ***X***_*i*_ and ***X***_*j*_ represent the coordinates of two cells. The repulsive Hookean force 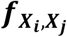 of stiffness ℱ acting on element ***X***_*i*_ is given by:

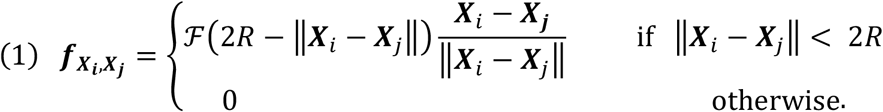

Since each cell can be in close proximity to several other cells, the repulsive force ***f***_*i*_ acting on that tumor cell combines contributions from *N*_*T*_ nearby tumor cells, and is given by the following equation:

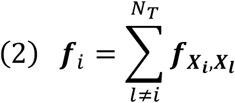

In addition, if the cell is migratory, a motility force ***g***_*i*_ will also act on that cell. The motility force can have a persistent direction or may dependent on the cues sensed from the microenvironment, such as contact guidance from the ECM fibers described below.

To resolve the overlapping conditions that may occur during cell division of migration, the tumor cells are relocated following the overdamped spring equation, where *ν* is the viscosity of the surrounding medium:

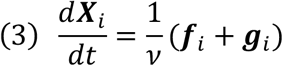

The ECM is modeled as a unit vector field ***h***(*x, y*) providing directions of the ECM fibers and scalar value *ξ*(*x, y*) representing the ECM stiffness. Initially, all fibrils have random directions. Subsequently, they can be realigned by the nearby cells. However, the extend of ECM remodeling depends on a combination of fibril stiffness and compliance to the direction of the moving or growing cells. This is described by the following equation:

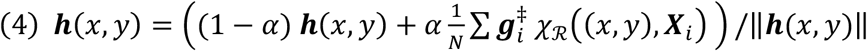

where, *α* is the compliance coefficient, *N* is the number of cells located within the neighborhood *χ*_ℛ_ of radius ℛ from the fibril, and 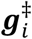 defines either a vector parallel to the direction of a migrating cell 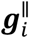 or perpendicular to the direction of a growing cell 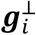.

The neighborhood relation is defined as follows:

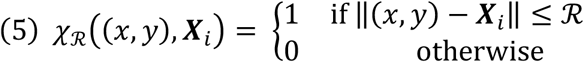

Fibril stiffness *ξ*(*x, y*) can increase at a rate Δ*ξ* due to pressure induced by active tumor cells, that is the cells that are either migrating or are relocated during tumor growth:

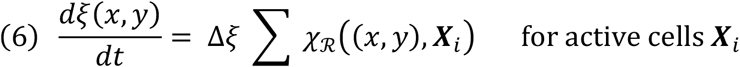

The motility force ***g***_*i*_ defining migration of the tumor cell ***X***_*i*_ is a consequence of a competition between the direction of the persistent cell movement 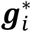 and the direction induced by contact-guidance ***h*** from *N*_*i*_ fibrils located in cell’s close neighborhood (*χ*_ℛ_). This is given by the following equation:

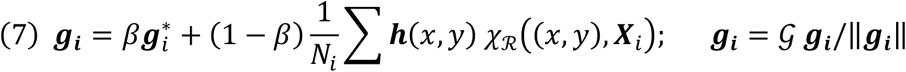

where *β* is a persistence coefficient and *𝒢* is the cell speed.

Model parameterization is based on our experimental measurements and scientific literature. We used tumor cell radius of *R* = 9.5*µm* (Figure 1 and (21)), repulsive force stiffness of ℱ = 0.35*µg*/*µm* · *s*^2^ (22), and medium viscosity of *ν* = 10*µg*/*µm* · *s* (23). The ECM grid width of *h* = 4*µm* represents a bundle of fibrils (Figure 1). Initially, we consider a uniform stiffness of *ξ*_0_ = 1 *σg*/*µm* · *s*^2^ (with scaling coefficient of *σ* = 10^−5^) that is characteristic for a normal mammary tissue (24), and can increase by Δ*ξ* = 0.02 *σg*/*µm* · *s*^2^ per a time step of Δ*t* = 15*s* to reach cancerous stiffness of *ξ*_*max*_ = 5*kPa* (24). The range at which the tumor cells and ECM fibrils can interact is assumed to be two fibril bundles wide, ℛ = 2*h*. The persistent direction of cell migration 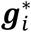, cell migration speed *𝒢*, as well as values of the persistent migration coefficient *β*, and the compliance coefficient *α*, will be varied in the examples discussed below.

## 3. Simulation Results

We used an off-lattice hybrid agent-based model (*MultiCell-LF*) to identify the rules of cell-ECM interactions that guide the development of various patterns of alignment of the ECM fibrils. We simulated migration of a single tumor cell through the ECM in section 3.1, and three diverse tumor associated collagen signatures: TACS-1 in section 3.2, TACS-2 in section 3.3., and TACS-3 in section 3.4 that are related to tumor behavior, such as benign growth or invasive migration.

### 3.1. ECM remodeling by a single migrating cell

Migration of a single cell through the ECM structure is used here to illustrate interactions between the cell and the nearby ECM fibrils. The direction of persistent migration was chosen to be 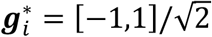, so the cell can move towards the upper-left corner of the domain with cell migration speed *𝒢* = 1*µm*/*s*. However, the actual cell movement is also influenced by the orientation of fibrils in the cell neighborhood *χ*_ℛ_, according to Eq.(7). The relation between cell persistent movement and movement due to contact-guidance from the surrounding fibrils is defined by the persistent migration coefficient *β*. In two examples presented in Figure 2, the cell moves either following the fibril orientations (Figure 2A, with the persistent migration coefficient *β* = 0.1) or ignoring the fibril orientation (Figure 2B, with persistent migration coefficient *β* = 0.9). Additionally, the migrating cell can modulate orientation of the nearby fibrils if they are compliant, according to Eq.(4). The two examples presented in Figure 2 show cases of either ECM with relatively stiff fibrils and moderate remodeling capabilities (Figure 2A, compliance coefficient *α* = 0.4) or ECM with high compliance to remodel which results in uniform orientation of all fibers behind the moving cell (Figure 2B, compliance coefficient *α* = 0.9). Thus, taking together, Figure 2 presents one case where the cell path towards the upper-left corner is tortuous because it follows the fibers with random orientations that remain relatively unchanged. The second case shows a cell that moves straight to the corner in a persistent fashion and also leaves behind a highly remodeled ECM. In these illustrative examples, we assumed that once the ECM is remodeled it will retain its new orientation and stiffness, so these effects can be visible at the end of simulation. In reality, there may be some elasticity effects that will make ECM fibrils to return to their initial configuration. This is not modeled here.

### 3.2. Formation of TACS-1

A normal breast tissue comprises of a branched structure of the mammary gland trees surrounded by a loose stromal connective tissue (Figure 3B and (25-27)). These ECM properties are mathematically modeled as a unit vector field of uniform low stiffness *ξ* = 1 *σg*/*µm* · *s*^2^ and random orientation ***h*** = (2 * [*rand, rand*] − 1)/‖***h***‖ (Figure 3A). This is an initial configuration for all our simulations and such fibril patterns are also observed experimentally in locations far from the growing cell colonies due to lack of interactions with tumor cells (see Figures 4-7 away from tumor cells). The TACS-1 signature is manifested by locally increased collagen density deposited by activated stromal cells (9, 11). While fibril orientation will not be changed in our model of TACS-1, the value defining local ECM density will be elevated.

### 3.3. Formation of TACS-2

During the tumor growth the expanding cell colony can impose pressure on the surrounding fibrils leading to changes in ECM alignment and stiffness. As a result, the elongated and straightened collagen fibrils were observed experimentally to encapsulate the tumor cluster (Figure 4B and (27, 28)). This is mathematically modeled as a change in ECM fibril orientation to be perpendicular to cell drag relocation force resulting in ECM alignment along the cluster boundary (Figure 4A). In this case the compliance coefficient *α* in Eq.(4) is small and the cell drag force 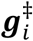 in Eq.(4) is perpendicular to the overall cell-cell interactions force acting on that cell, i.e.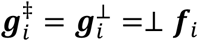. Moreover, every time that the fibril orientation is changed due to a push from the growing cell cluster, the stiffness of the fibril bundle at the point of contact with a cell is increased, according to Eq.(6). This is illustrated in Figure 5.

An individual cell (Figure 5A) is located in the non-rigid randomly oriented ECM fibril structure. Upon division, both daughter cells push on one another to resolve cell overlapping and, in the process, they push on the surrounding ECM fibrils. The fibrils exert resistance and realign perpendicular to the pushing cells increasing also their stiffness (Figure 5B). This process is elevated when the cell cluster grows in size and more cells are pushing on the nearby fibrils (Figure 5C-E).

### 3.4. Formation of TACS-3

During the emergence of tumor cell invasive cohorts, the ECM fibrils are primarily aligned in the direction of cell migration and are perpendicular to the tumor boundary (Figure 6B and (10, 27)). The algorithm to achieve that in our model is to change orientation of ECM fibrils to parallel to cell motility force and increase fibril stiffness at the point of contact. In this case, the compliance coefficient *α* in Eq.(4) is small and the cell drag force 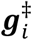 in Eq.(4) is parallel to the direction of the persistent cell movement, i.e. 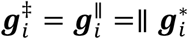. Here, the persistent coefficient is large (*β* = 0.99) and the cells ignore directions of the nearby fibrils. The direction of cell persistent migration was chosen to be 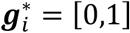, so the cells move vertically with cell migration speed *𝒢* = 1*µm*/*s*. In addition, every time that the fibril orientation is changed due to the pull from the migrating cell cluster, the stiffness of that fibril bundle is increased, according to Eq.(6). This is illustrated in Figure 7.

**Figure 6.**
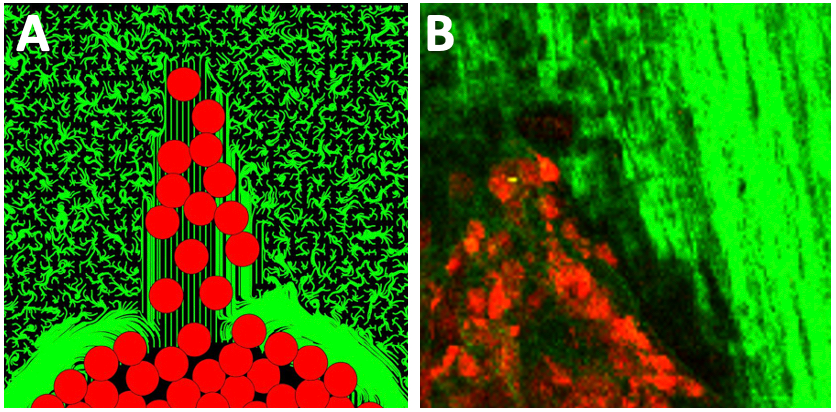
Model of TACS-3. **A**. Simulated ECM with fibrils aligned radially out of the growing cell cluster at the side of cell invasions and **B**. the corresponding ECM pattern around the invading cells from intravital fluorescent microscopy.

**Figure 7.**
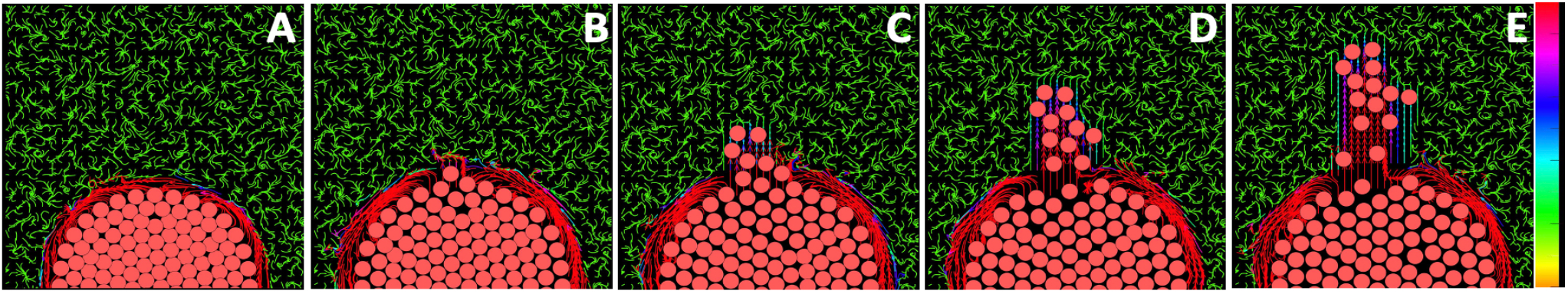
Development of TACS-3 in the mathematical model. **A-E**. Snapshots from a simulation showing the emergence of stiff ECM fibrils aligned parallel to the migrating cell cohort. The local ECM fibril stiffness is shown in a color scale.

Initially, the cluster of cells resides in the ECM that has a random orientation everywhere except of the area near the tumor boundary (Figure 7A). Upon initiation of the invasion process, a single tumor cell starts migrating vertically and remodeling the nearby ECM fibrils by aligning them in the same direction (Figure 7B). Subsequently the cells move in the same vertical direction and exert pulling forces on the nearby fibrils living behind a wide band of vertically aligned fibrils (Figure 7C-D).

## 4. Discussion

In this paper, we used an off-lattice hybrid agent-based Multi-Cellular Lattice-Free (*MultiCell-LF*) model to identify the rules of cell-ECM interactions that lead to the emergence of various patterns of alignment of the ECM fibrils. It has been observed experimentally and in clinical samples that three specific tumor-associated collagen signatures (TACS) are characteristic for three stages of cancer progression (5, 10, 11). The TACS-1 signature that has a wavy appearance with unorganized fibrils was detected in the areas located far from the growing tumor cell colony. The TACS-2 signature was characterized by ECM fibrils aligned parallel to the edge of the tumor cluster. Finally, the TACS-3 signature was described as ECM fibrils oriented radially from the tumor cluster. Using our mathematical model, we identified rules of cell-ECM interactions that resulted in the given fibril alignment. Our starting point was the ECM with uniformly low stiffness and random fibril orientation. For the TACS-1 signature, the fibril orientation was not changed, but the local ECM density was increased. For the TACS-2 pattern, the fibril orientation was changed to be perpendicular to the force from a growing cell cluster and stiffness of the fibril bundle was increased at the point of contact with the cells. For the TACS-3 pattern, the fibril orientation was modified to be parallel to the direction of the persistent cell migration and stiffness of the fibril bundles in contact with the moving cells was increased.

While we were able to identify cell-ECM interactions that resulted in the three TACS signatures, there are still some open questions that can be addressed in the future applications of the model that we have developed. It is not known what processes can lead from one signature to the other. The experimental observations were made using fixed tissue samples that do not allow to trace the signature progression. The mathematical modeling can provide a way to test various hypotheses of signature evolution. It is also not known if the emergence of TACS-3 signature precedes tumor cell invasion, so the cells are utilizing the already existing fibril tracts to migrate, or if TACS-3 is the consequence of cell invasion, so the migrating cells leave the fibril tracks behind during their movement? Being able to identify the rules of transition from one TACS signature to the other may help in the future in cancer diagnosis, in prognosis of tumor progression, and may serve as histology-based biomarkers.

